# Loop-extrusion and polymer phase-separation can co-exist at the single-molecule level to shape chromatin folding

**DOI:** 10.1101/2021.11.02.466589

**Authors:** Mattia Conte, Ehsan Irani, Andrea M. Chiariello, Alex Abraham, Simona Bianco, Andrea Esposito, Mario Nicodemi

## Abstract

Loop-extrusion and phase-separation have been proposed as mechanisms that shape chromosome large-scale spatial organization. It is unclear, however, how they perform relative to each other in explaining chromatin architecture data and whether they compete or co-exist at the single-molecule level. Here, we compare models of polymer physics based on loop-extrusion and phase-separation, as well as models where both mechanisms act simultaneously in a single molecule, against multiplexed FISH data available in human loci in IMR90 and HCT116 cells. We find that the different models recapitulate bulk Hi-C and average microscopy data. Single-molecule chromatin conformations are also well captured, especially by phase-separation based models that better reflect the experimentally reported segregation in globules of the considered genomic loci and their cell-to-cell structural variability. Such a variability is consistent with two main concurrent causes: single-cell epigenetic heterogeneity and an intrinsic thermodynamic conformational degeneracy of folding. Overall, the model combining loop-extrusion and polymer phase-separation provides a very good description of the data, particularly higher-order contacts, showing that the two mechanisms can co-exist in shaping chromatin architecture in single cells.

## INTRODUCTION

To understand the machinery that in the nucleus of cells establishes at large scales the 3-dimensional (3D) architecture of chromatin^1–14^, encompassing DNA loops^15^, Topologically Associated Domains (TADs)^16,17^ and other structures^13,18^, different physical mechanisms have been proposed and investigated via models relying solely on fundamental physical processes^19–45^ and via computational approaches^46–59^. However, it remains unclear how well different mechanisms capture folding at the single molecule level, how they compare against each other in explaining experimental data and whether they compete or co-exist in determining the structure of chromosomes. Here, we explore two recently discussed classes of models that focus on two distinct physical mechanisms, respectively loop-extrusion and polymer phase-separation, that we compare against single-molecule super-resolution microscopy^6^ and bulk Hi-C data^15,60^ available in human loci in IMR90 and HCT116 cells.

Loop-extrusion and phase-separation based polymer models of chromosomes reflect two classical, yet distinct scenarios of molecular biology to explain the formation of DNA contacts^61^. The first class considers the picture where physical proximity between distal sites is established by molecular motors that bind to DNA and extrude a loop^19,20,31,39,40^. This is an out-of-equilibrium, active physical process that involves energy, e.g., ATP, consumption. The model envisages that those loop-extruding complexes stochastically bind to a polymer chain and extrude loops until encountering another motor, an anchor site or unbinding from the chain. While the polymer becomes compacted in a linear array of loops, specific contacts are established between the motor anchor sites where extrusion halts, hence defining boundaries between subsequent chromatin regions. Experimental evidence indicates that Cohesin and Condensin can be components of the motor complex, while properly oriented CTCF sites can act as anchor points^40^. Computer simulations have shown that such a model can explain with good accuracy loops and TADs visible in bulk Hi-C contact maps in interphase as well as, for example, in mitotic chromosomes^19,20,31,39,40^. Variants of such a model have been also developed where chromatin loops are formed by thermal random sliding of DNA into an extruding molecule^31^ or by, e.g., transcription-induced supercoiling^39^.

The second class of polymer models^21–30,32–38,41–45^ considers another classical scenario where physical proximity between distal DNA sites results from interactions mediated, for instance, by diffusing cognate bridging molecules, such as Transcription Factors, or from direct interactions produced, e.g., by DNA bound histone molecules. In the Strings and Binders (SBS) model^42,44^, for example, a chromatin filament is represented as a self-avoiding chain of beads, along which are located different types of binding sites for cognate diffusing binders that can bridge those sites. The binding sites have been correlated to different molecular and epigenetic factors, ranging from active and poised Pol-II to eu- and heterochromatin sites^21,27,28,45^. The steady-state 3D conformations of the system are determined by the laws of physics and fall in different structural classes corresponding to its thermodynamics phases. In the SBS model, for instance, upon increasing the concentration or affinity of binders, the system undergoes a polymer phase-separation transition from a coil, i.e., randomly folded, to a globular state, where distinct globules self-assemble along the chain by the interactions of cognate binding sites^24,27,35,44^. Polymer physics explains that thermodynamic phases are independent of the specific origin of the interactions - e.g., direct or mediated by diffusing factors - so different models can belong to the same universality class^62^. For that reason, the thermodynamic phases of, say, the SBS model also occur in models with direct chromatin interactions. Those phase transitions result in structural changes of the chain that spontaneously establish contact or segregation of specific, distal sites, such as genes and their regulators. Such a class of models has been shown to explain Hi-C, SPRITE, GAM and microscopy contact data across the genome, from the sub-TAD to chromosomal scales^21–30,32– 38,41–45^, also at the single molecule level^35,38^.

It is unclear, however, how loop-extrusion and polymer phase-separation perform relative to each other in capturing chromatin folding and whether they compete or co-exist in establishing chromosome architecture. Here, we implemented different versions of those models to benchmark their structural predictions at the single-molecule level against independent multiplexed FISH data^6^. We simulated first a simple loop-extrusion (LE) model^20^ of the considered loci. Next, we developed an extended LE (eLE) model whose anchor site genomic locations are optimized to best fit experimental contact data. Additionally, to mimic epigenetic differences of single cells, in the model the anchor sites can differ across single molecules^29^. We also considered the SBS model of the studied loci^35^ and, finally, we introduced a model combining eLE and SBS (the LE+SBS model), i.e., a model where in a single molecule both LE and SBS mechanisms act simultaneously. We find that both loop-extrusion and phase-separation based models can explain well ensemble-averaged microscopy and bulk Hi-C data, albeit the simple LE model is only partially effective. Our single-molecule analyses show that both types of models do capture the main features of single-cell chromatin conformations and higher-order contacts. Yet, phase-separation based models better reflect the experimentally reported segregation in globules of the considered genomic loci and their cell-to-cell structural variability. Such a variability results from two main concurrent sources: the intrinsic thermodynamic degeneracy of polymer folding and single-cell epigenetic heterogeneity. Consistent with such a picture, the LE+SBS model turns out to provide overall an excellent description of all the different datasets and to have the least discrepancy with microscopy triple contact data, supporting the view that loop-extrusion and phase-separation can co-exist at the single-molecule level in determining chromatin architecture.

## RESULTS

### Polymer models of the studied loci

We implemented the polymer models of two 2Mb wide loci in human IMR90 and HCT116 cells where, as stated, single-cell super-resolution microscopy data^6^ are available at 30kb resolution (**Fig. 1a, Suppl. Fig. 1a**). To assess the role of the different ingredients of the models, we developed distinct versions that we compared against single-cell data.

**Figure 1.**
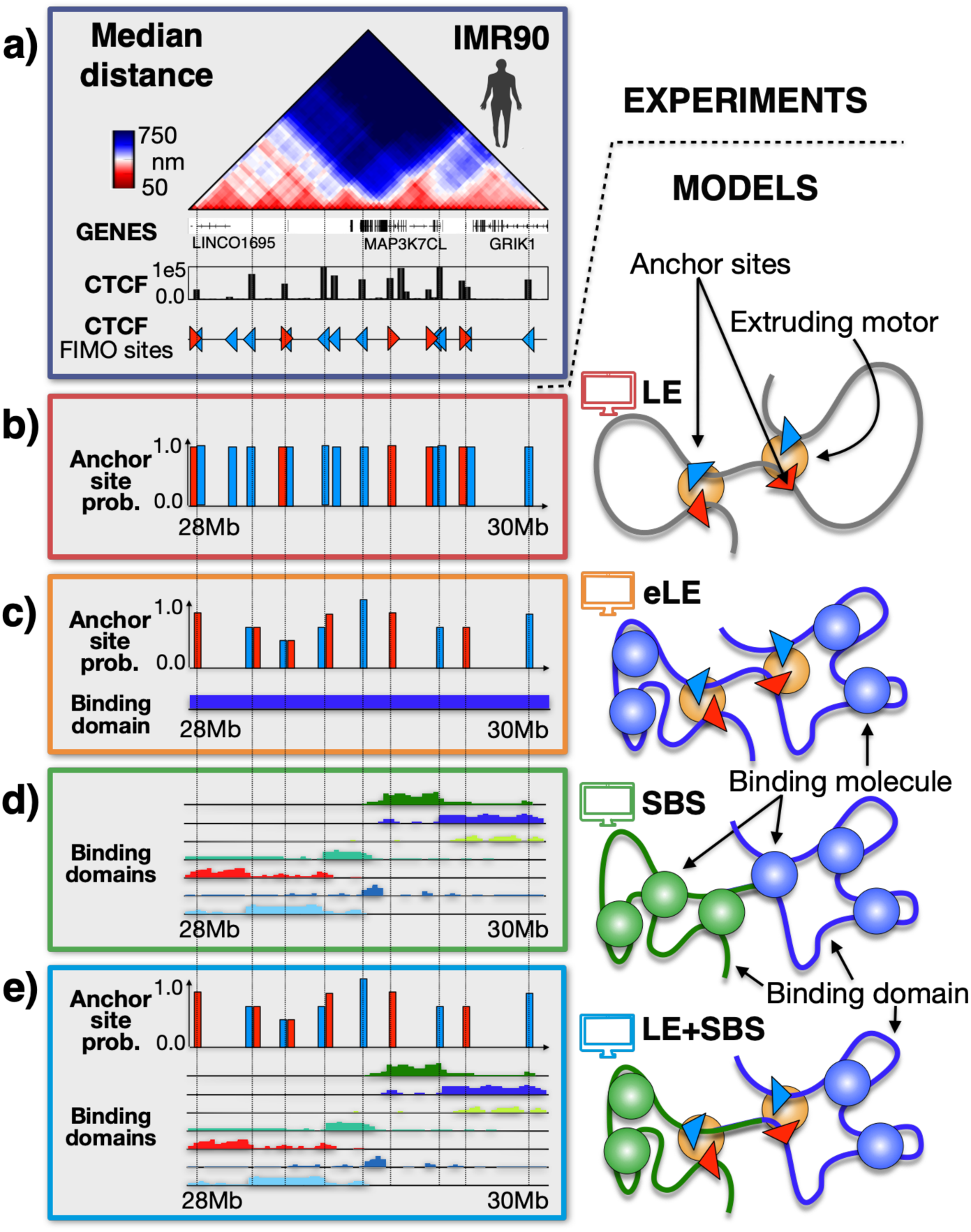
Scheme of the investigated polymer models. We used Molecular Dynamics simulations to investigate polymer models where folding is based on two different physical processes: (i) DNA loop-extrusion and (ii) polymer phase-separation, recapitulated respectively by the LE^19,20^ and by the SBS models^42,44^. **a)** Microscopy median distance^6^ and ENCODE^64^ CTCF data are shown for the studied 2Mb wide locus in human IMR90 cells. **b)** We considered a simple Loop-Extrusion (LE) model^20^ where active motors extrude polymer loops until encountering another motor or CTCF anchor points with opposite orientation, which are fixed and equal in all single-molecule simulations. **c)** We also considered an extended version of the LE (eLE) whose anchor site locations are optimized, independently of CTCF, to best reproduce Hi-C and average microscopy data. To represent the epigenetic heterogeneity of single cells, those anchor sites have a finite probability to be present in a model single molecule^29^. **d)** In the Strings and Binders (SBS) model^35^ a chromatin filament is represented as a self-avoiding chain of beads including different types of binding sites (colors) for diffusing cognate binders that can bridge those sites, hence driving a micro-phase-separation of the chain in distinct globules. The binding site locations are determined by the PRISMR method and correlate with different combinations of epigenetic factors including, but not limited to, CTCF and cohesin^28,35^. **e)** We also considered a polymer model (LE+SBS) where in a single molecule both the eLE and SBS mechanisms act simultaneously.

First, we implemented a simple LE model^20^, where loop-extruding motors stochastically bind to a polymer bead chain and extrude loops until encountering anchor points with opposite orientation or another motor or unbinding from the chain (**Fig. 1b** and **Methods**). The position and orientation of the anchor points are identified by the FIMO standard motif finding analysis^63^ based on the peaks of CTCF ChIP-seq data from ENCODE^64^. While the motors can stochastically bind to and unbind from the chain, the anchor sites are fixed and equal in all single-molecule computer simulations. Their anchoring strength is set to 100%, i.e., when an extruder arrives at an anchor point it remains blocked at that position, yet we checked that the overall results do not change for strengths in the range down to 60% (**Methods**). This model is hereafter referred to as the LE model. To explore the potential of the loop-extrusion mechanism beyond such a minimal implementation, we also considered a more refined version where, to mimic epigenetic differences across single cells, each anchor site is present, with a given probability, only in a subset of model single-molecules^29^ (**Fig. 1c, Suppl. Fig. 1b** and **Methods**). Additionally, to best reproduce population-averaged Hi-C and microscopy distance data, we searched for the optimal genomic location and probability of the motor anchor sites, independently of CTCF tracks (**Methods**). In the considered loci, most of those optimal sites coincide with FIMO CTCF peaks (**Fig. 1c, Suppl. Fig. 1b**), but not all, and conversely many FIMO CTCF peaks are not included as model anchor sites. The probability to be present in a model single-molecule is found to be different for different anchor sites, ranging from roughly 50% to 100% (**Fig. 1c, Suppl. Fig. 1b**), values consistent with current estimates of cell epigenetic heterogeneity^65^. Finally, to better fit the features of Hi-C and microscopy data, such as TADs and globules (see below), the beads of the polymer chain are subject to a self-interaction produced by unspecific bridging molecules. Such a variant of the LE model is hereafter named the extended LE (in short, eLE).

Next, in the considered loci we implemented the SBS model^35^ whereby chromatin is represented as a self-avoiding chain of beads, in a thermal bath, with specific binding sites for cognate diffusing molecular binders (**Fig. 1d, Suppl. Fig. 1c** and **Methods**). The location and types of binding sites are different for the different loci and are inferred via a machine learning procedure based on the PRISMR method, which takes as input only Hi-C data^28,35^. The model of the HCT116 locus has four binding site types and the model of the IMR90 locus has seven types, each visually represented by a different color along the chain (**Fig. 1d, Suppl. Fig. 1c**). The binding site types have been shown to correlate with different combinations of chromatin architectural factors, such as CTCF/Cohesin, H3K4me3 or H3K4me1^35^. As mentioned above, the equilibrium 3D conformations of the SBS model fall in structural classes corresponding to its thermodynamics phases^62^: upon increasing binder concentration or affinity above a threshold value, the system undergoes a phase transition from a coil (i.e., randomly folded) to a polymer phase-separated state where distinct, compact globules self-assemble along the chain in correspondence of its different, prevailing binding domains (i.e., locally enriched colors)^35^. The intrinsic thermodynamic degeneracy of the states of the model results in a broad variety of 3D single-molecule conformations^35^ (**Methods**). We also developed a variant of the SBS model where cognate DNA sites have direct physical interactions, rather than mediated by binders, and our overall findings remain unchanged (**Suppl. Fig. 2, Methods**) as expected from Statistical Mechanics^62^.

Finally, to check whether active mechanisms, such as loop-extrusion, and passive mechanisms, such as thermodynamic polymer phase-separation, could coexist to shape chromatin architecture in the studied loci, we implemented a polymer model combining the above described eLE and SBS models, i.e., a model where both mechanisms act simultaneously in each single molecule (named the LE+SBS model, **Fig. 1e, Suppl. Fig. 1d** and **Methods**). For each of the considered models, an ensemble of 3D conformations was obtained via Molecular Dynamics simulations in the steady state^29,35^ (**Methods**). In all the considered cases the model unit length scale was mapped into physical units by equating the median gyration radius to its corresponding experimental counterpart^6,35^ (**Suppl. Fig. 3, Methods**).

### Both loop-extrusion and phase-separation based models recapitulate average microscopy and Hi-C data

To benchmark the different models, we focused first on how they recapitulate population-averaged experimental data by comparing their median distance and contact maps against, respectively, multiplexed FISH^6^ and bulk Hi-C data^15,60^.

In our IMR90 case study locus, we found that the models all capture the global patterns visible in the median distance matrix^6^ (**Fig. 2a, Methods**). To have a quantitative measure of similarity, we computed the genomic distance-corrected Pearson correlation coefficient, r’, between model and experiment. The LE has the lowest r’ (r’=0.19), while the eLE has r’=0.49, highlighting a markedly improved similarity to the experiment. The data appear to be better captured by the SBS and by the LE+SBS models, as signaled by their higher correlations (r’=0.77 and r’=0.70, respectively). Analogous results are found by comparing the model contact matrices against Hi-C data^15^ (**Fig. 2b, Methods**): LE has the lowest correlation (r’=0.24), eLE has r’=0.57, while SBS and LE+SBS models comparatively better reproduce Hi-C contact patterns (r’=0.74 and r’=0.72, respectively). We also considered other measures of similarity, such as the simple Pearson correlation (**Suppl. Table I**), which provided analogous results.

**Figure 2.**
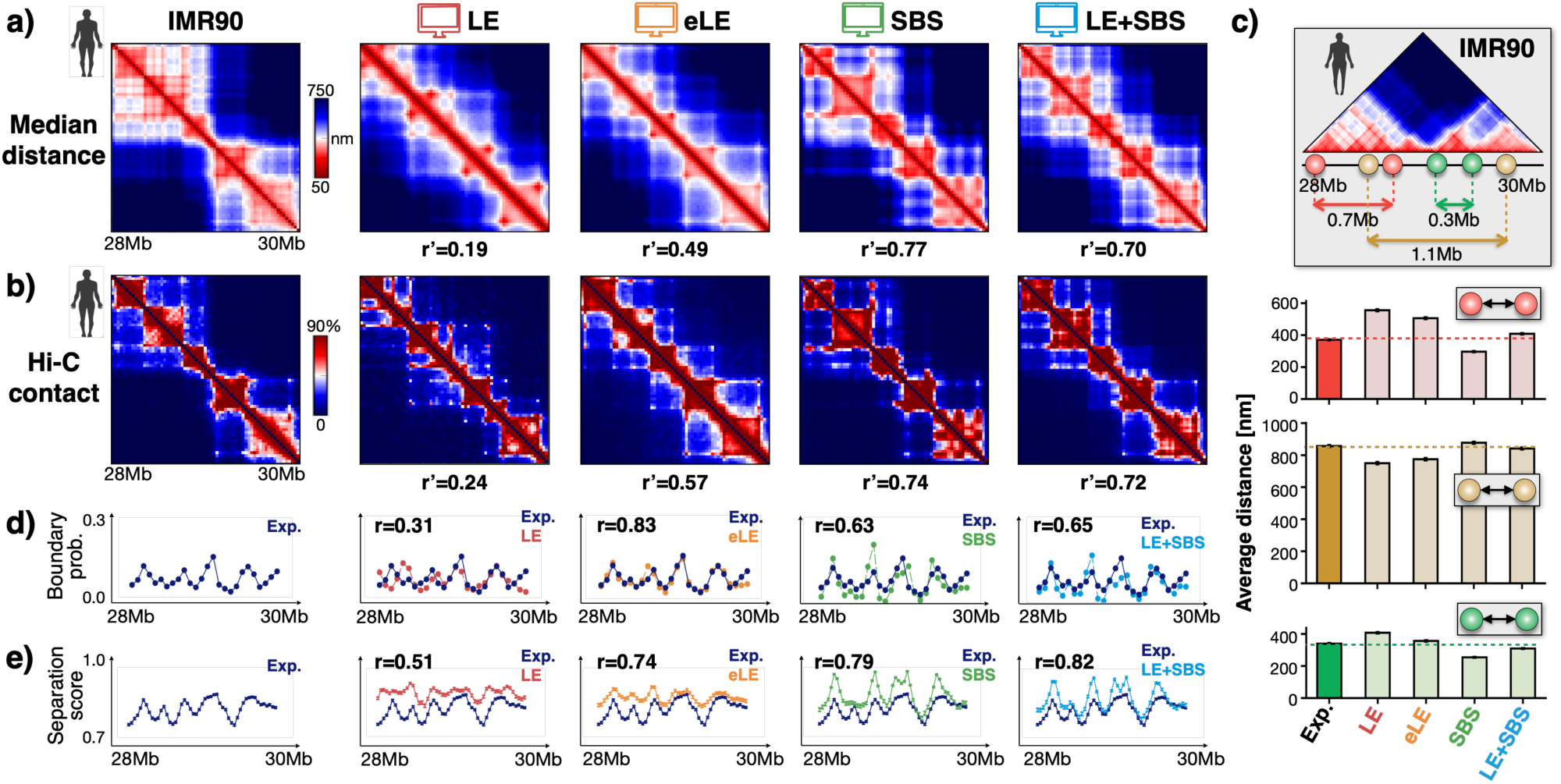
Both loop-extrusion and phase-separation based models recapitulate bulk Hi-C and average microscopy data. **a)** In-silico median distance and **b)** average contact data are compared to microscopy^6^ and Hi-C^15^ data (left) in the IMR90 locus. The different models have high genomic distance-corrected Pearson correlations (r’) with the experiments, particularly the eLE, SBS and LE+SBS models. **c)** The model derived average distances are reported for specific pairs of sites separated by TAD boundaries (yellow) or connected in loops within a TAD (green) or across a TAD boundary (red). **d)** The average single-molecule genomic boundary probability and **e)** the separation score are also well recapitulated by the models.

Next, we focused on the relative distances of specific, interesting pairs of sites in the IMR90 locus (**Suppl. Table II**). We considered: (i) a pair of sites (green, **Fig. 2c** and **Suppl. Table II**) located 0.3Mb apart from each other within the same TAD, having a strong interaction; (ii) a pair of sites (red), located 0.7Mb away in different sub-TADs, having a strong loop contact in the median distance matrix; and (iii) a pair of 1.1Mb distant sites (yellow) from different TADs, separated by a strong TAD boundary. Albeit the genomic separation of the red pair is twice as large than the separation of the green, those pairs have a similar average distance in the experiment, close to 400nm, whereas the boundary separated yellow pair is more than 800nm apart (**Fig. 2c, Suppl. Table II**). We found that the different models all recapitulate those values (**Fig. 2c, Suppl. Table II**) and, interestingly, the LE+SBS model is overall the closest to the experiment across those specific pairs of sites. Additionally, we checked that the distance distributions derived from the models are all similar to the corresponding microscopy distance distributions (**Suppl. Fig. 4**). We stress, however, that the specific values of those distances can depend on the minute details of the models, such as the shape of the interaction potential or loop-extruder size, so the agreement could be further improved.

To assess how well distinct models capture different aspects of chromatin folding, we also computed the probability to find a TAD boundary at a given genomic location and the average separation score^6^ along the locus in single-molecule conformations (**Fig. 2d**,**e, Methods**). In the IMR90 case study locus, we found that the boundary probability and the boundary strength averaged over all genomic positions are similar across the different models and very close to the experimental values (**Suppl. Fig. 5**). The boundary probability as a function of the genomic coordinates of the locus, however, is better captured by the eLE model, which has the highest Pearson correlation with experimental data (r=0.83, **Fig. 2d**), while the LE has the lowest correlation (r=0.31). The SBS and LE+SBS models also provide a good fit to the data, having respectively r=0.63 and r=0.65. We also found that all the models provide a good overall description of the average separation score along the locus (**Fig. 2e**): the LE has the lowest correlation to the corresponding experimental data (r=0.51), the eLE has r=0.74, the SBS r=0.79 and the LE+SBS model r=0.82.

Our analysis of the HCT116 locus returned a very similar picture about the performance of the different models to describe average distance and Hi-C data (**Suppl. Fig. 6a, b**) as well as TAD boundary probabilities and separation scores (**Suppl. Fig. 6c, d and Suppl. Fig. 7**).

Taken together, our results show that both loop-extrusion and phase-separation based models are consistent with ensemble-averaged microscopy and bulk Hi-C data. While the LE model is only partially effective, the eLE, which incorporates a single-molecule variability of optimized anchor sites, works well in the description of those data and is the best to recapitulate the TAD boundary probability function. However, polymer models including globule phase-separation mechanisms (SBS and LE+SBS) have overall higher correlation values with average microscopy distance and Hi-C contact data, and better capture some local features of chromatin folding, such as the separation score.

### The models are overall consistent with chromatin structure at the single-molecule level

To compare how the different models describe chromatin structure at the single-molecule level we took advantage of the mentioned super-resolution microscopy data^6^ and of the ensemble of polymer 3D conformations produced via our computer simulations.

First, we checked how well each model represents single-cell chromatin conformations by performing an all-against-all comparison of single-molecule imaged and model 3D structures. We used a method^35,66^ whereby each 3D conformation from microscopy data is univocally associated to a corresponding model structure (for each considered type of model) by searching for the least root mean square deviation (RMSD) of their coordinates (**Fig. 3a, Suppl. Fig. 10a and Methods**). To test the statistical significance of the association, we compared the RMSD distribution of the best-matching experiment-model pairs against a simple control case where the RMSD distribution is computed between random pairs of imaged structures. We verified that for each of the considered polymer models the RMSD distribution of the best-matching pairs is statistically different from the control in both the IMR90 and HCT116 loci (**Fig. 3b** and **Suppl. Fig. 8a**, two-sided Mann–Whitney test p-value = 0). Quantitively, in the IMR90 locus we found, consistently across the models, that less than 5% of the former distribution is above the first decile of the control (**Fig. 3c**) and, in particular, the SBS model performs slightly better than the others. The analysis of the models of the HCT116 locus returned similar results (**Suppl. Fig. 8b**). As an additional test, we also considered a more stringent control where the RMSD is computed only between pairs of imaged structures having overall similar distance matrices, i.e., with a corresponding genomic distance-corrected correlation larger than 0.5 (i.e., with r’>0.5, see below), and we found analogous results (**Suppl. Fig. 9**). Hence, the model conformations best matching the experimental structures have a statistically significant RMSD distribution and provide a non-trivial description of chromatin molecules in single cells (**Fig. 3a, Suppl. Fig. 10a**).

**Figure 3.**
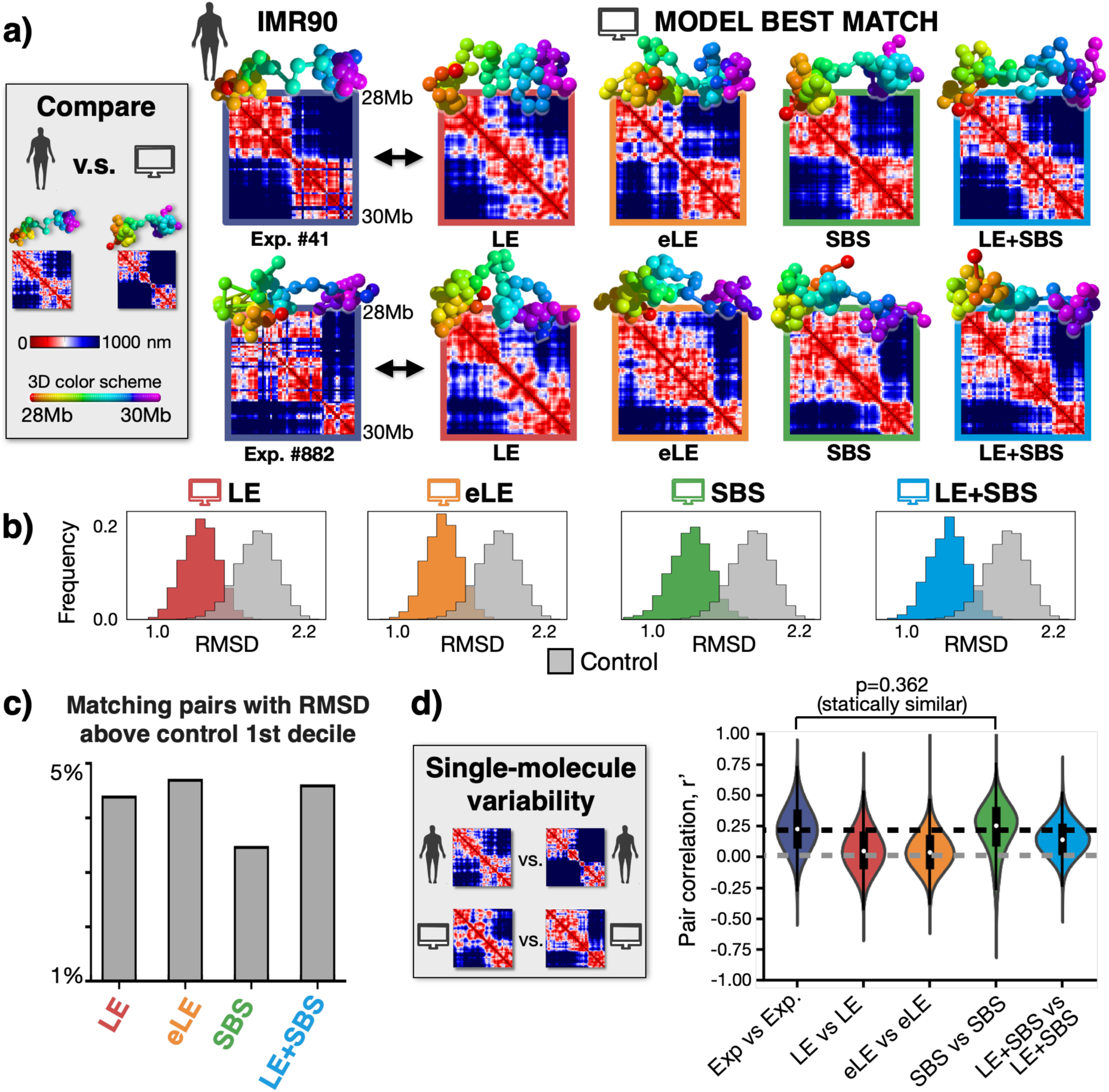
Single-cell chromatin conformations are well captured by the models, especially by phase-separation based ones. **a)** Microscopy single-cell chromatin structures of the IMR90 locus^6^ (left) are associated to a best matching single-molecule conformation in each model via the minimum RMSD criterion. Here two examples are shown. **b**) For each of the considered polymer models, the RMSD distribution of the best-matching experiment-model pairs is statistically different from a control RMSD distribution made of random pairs of experimental structures (two-sided Mann–Whitney test p-value = 0). **c)** Less than 5% of best matching pairs have an RMSD above the 1st decile of the control distribution. **d)** The variability of microscopy single-molecule structures is measured by the distribution of r’ correlations between pairs of distance matrices and is compared to the variability of in-silico structures. The r’ distribution of the SBS model is statistically indistinguishable from the experimental one (two-sided Mann–Whitney test p-value = 0.362).

Next, we tested whether the variability of the ensemble of model single-molecule structures reflects the experimentally observed variability of chromatin single-cell conformations^6^. In the IMR90 locus, for example, the distribution of r’ correlations between pairs of experimental single structure distance matrices has an average r’=0.23 and a variance equal to 0.18 (**Fig. 3d**), showing that while the imaged structures are broadly varying they have also a significant degree of similarity^6,35^. For each model, we computed the corresponding distribution of r’ correlations between all model single-molecule distance matrices and we compared it with the experimental one (**Fig. 3d** and **Methods**). Interestingly, the r’ distributions of the different models have all a shape similar to the experiment and a similar variance, yet they have different average values (**Fig. 3d**). The LE and eLE model average r’ (r’=0.06 and r’=0.04 respectively) is significantly lower than the experimental value, showing that their single-molecule structures have a lower degree of similarity with each other than single-cell imaged chromatin conformations. The LE+SBS model has an average r’=0.14, while the SBS model has r’=0.23, which is equal to the microscopy value (**Fig. 3d**). In fact, the r’ distribution of the SBS model is statistically indistinguishable from the experimental distribution (two-sided Mann–Whitney test p value = 0.362), while the other models are statistically different (p <0.001). Additionally, we verified that analogous results are found if the experiment-experiment r’ distribution is compared to the distribution of r’ correlations between experiment and model single-molecule distance matrices (**Suppl. Fig. 11a**). We stress, again, that those correlation measures can depend on the minute details employed to construct the models and the agreement with the experiment could be further improved. Finally, the analysis of the HCT116 locus returns very similar results to those of the IMR90 locus (**Suppl. Fig**.**s 10b, 11b**).

In summary, consistent with our findings on bulk data, our single-molecule analyses support the view that the different polymer models all provide a non-trivial description of single-cell chromatin conformations. While both loop-extrusion and phase-separation based models capture the main features of chromatin single-molecules, in the studied loci we find that the latter models better reflect the microscopy observed single-molecule globular structure and variability. In particular, our analysis shows that chromatin structure variability across single cells results from two main distinct, yet concurrent sources: on the one hand from the intrinsic degeneracy of folding that we find in all the considered models, and on the other hand from the differences of anchoring points (or, analogously, binding sites) in single-molecules, representing the epigenetic heterogeneity of single cells.

### The models well reproduce microscopy triple contact data

To assess how well the different models capture higher-order contacts, we investigated their predicted average triplet contact probability matrix, which we compared to microscopy data^6^ (**Methods**). We focused on triplets formed by six different genomic viewpoints roughly equally spaced along the IMR90 locus that correspond to some main TAD boundaries and loops of the pairwise median distance matrix (**Fig. 4** and **Suppl. Fig. 12**). In our analysis, by definition, a triplet is formed if three genomic sites have all their pairwise distances below a threshold value. The triplet probability depends on such a threshold, but we checked that the measured values are proportionally conserved if the threshold is varied around 150nm in a range from 100 to 200nm (**Methods**).

**Figure 4.**
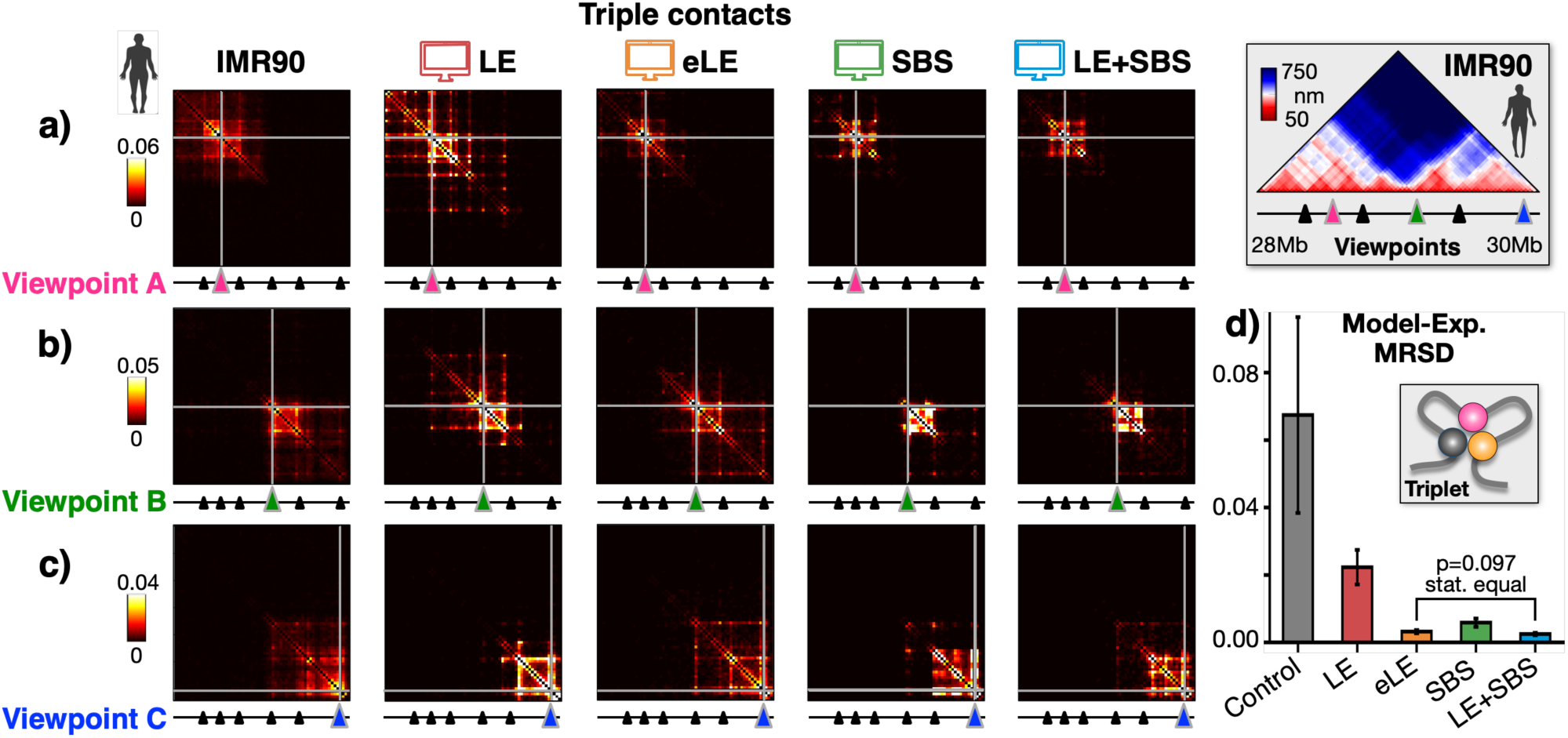
Triple contact data are well described by the models, especially by the eLE and the LE+SBS. Triple contact probability maps are shown in microscopy data^6^ (left) and in the models from three different viewpoints (**a), b), c)**, more viewpoints in **Suppl. Fig. 12**). **d)** The mean relative squared difference (MRSD) between imaging and model triplet contact maps is the lowest in the LE+SBS model, which is statistically equivalent to the eLE model (two-sided Welch’s t-test p=0.097). Their MRSDs are, instead, statistically different from both the LE, the SBS and the control (two-sided Welch’s t-test p<0.001). The control is made of randomly folded self-avoiding polymer chains with same number of beads and size than the experimental structures.

Microscopy data reveal that triplets are typically compartmentalized in the studied loci and restricted to the TAD encompassing each of the selected viewpoints (**Fig. 4a-c** and **Suppl. Fig. 12**), showing that TADs tend to create local environments where also multiple contacts become enriched. The different polymer models do capture experimental triplet patterns across all the considered viewpoints. To quantitatively assess the similarity between experiment and model predicted triplets, we computed the mean relative squared difference (MRSD) between the corresponding entries of the two matrices over the studied viewpoints (**Fig. 4d, Methods**). To set a reference, we also considered the triplets formed in a random control made of self-avoiding-walk (SAW) polymer chains having the same number of beads and gyration radius (i.e., linear size) as the real images of the locus (**Methods**). Our analysis shows that the LE model has an MRSD with the experiment that is one third of the random control value, yet it has the largest discrepancy with the experiment compared to the other considered models, whose MRSD is at least one order of magnitude smaller than the control. Interestingly, the LE+SBS model has the lowest distance from the experiment and its MRSD is statistically different from both the LE, the SBS and control case (**Fig. 4d**, two-sided Welch’s t-test p<0.001), whereas it is statistically equal to the eLE MRSD (two-sided Welch’s t-test p=0.097).

Taken together, our results show that both loop-extrusion and phase-separation mechanisms can explain higher-order contacts. However, a model combining both mechanisms (LE+SBS) turns out to have the least discrepancy with microscopy triplet data and overall provides an excellent description of all the different experimental datasets considered, supporting the view that loop-extrusion and phase-separation can co-exist in single-molecules in establishing chromatin architecture.

## DISCUSSION

To investigate the physical mechanisms that shape chromatin 3D large scale organization, we explored via Molecular Dynamics simulations two classes of polymer models where folding is based on two distinct physical processes: DNA loop-extrusion and polymer phase-separation, recapitulated respectively by the LE and by the SBS models (**Fig. 1**). We assessed how they perform relative to each other in capturing chromatin bulk Hi-C contact^15,60^ and single-molecule microscopy data^6^ in human IMR90 and HCT116 cells, and we exploited such data to establish whether those mechanisms compete or coexist in single cells.

We considered, first, a simple loop-extrusion (LE) model^20^ of those loci (**Fig. 1b**). Next, we introduced an extended version of the LE (named eLE, **Fig. 1c**), where the genomic locations of the extruding motor anchor sites are optimized, independently of CTCF peaks, to best reproduce population-averaged experimental data. Additionally, to mimic epigenetic differences among single cells, each of those anchor sites has a specific probability to be present in a model single molecule^29^. The probability values returned by the optimization search range from 50% to 100%, consistent with current estimates of cell epigenetic heterogeneity^65^. Interestingly, most anchor sites of the optimal eLE model are found to coincide with CTCF peaks (**Fig. 1c**), but not all, and conversely many CTCF peaks are not taken as anchor sites, hinting that CTCF may be combined with other signals in anchoring loop-extruding motors^67^. Considering the basic ingredients that inform the LE model, we find that it performs well to fit experimental data. Yet, the eLE model better recapitulates average microscopy and Hi-C data and higher order contacts in single-molecules.

We also considered the SBS model of the studied loci (**Fig. 1d**), i.e., a model where the attraction between cognate binding sites on the polymer chain and their associated binding molecules drives a micro-phase-separation of the chain in distinct globules^35^. For completeness, we checked that a model with direct interactions between binding sites (rather than mediated by diffusing binders) has behaviors analogous to the SBS. Finally, we introduced a model combining the molecular elements of the eLE and of the SBS (the LE+SBS model) where in a single molecule both the LE and SBS mechanisms act simultaneously (**Fig. 1e**). We find that the SBS and LE+SBS models explain well bulk Hi-C and single-molecule microscopy data, and reflect the experimentally reported chromatin segregation in globules and its cell-to-cell structural variability more accurately than the LE or eLE models.

Importantly, a further optimization of the model fine details, such as the employed specific interaction potentials (shape, depth, distance of the potential minimum, etc.) or the specific nature of the modelled DNA extruding motors (size, speed, directionality, etc.), can on one hand improve even more the model agreement with experiments and on the other hand provide additional mechanistic information. Nevertheless, the models here investigated perform well considering their simplicity (**Fig**.**s 2-4**). In particular, the LE+SBS model returns an overall excellent description of the different datasets and the least discrepancy with microscopy triplet data, showing that loop-extrusion and phase-separation can co-exist in shaping the complex chromatin architecture of the studied loci. Our analyses also illustrate that the experimentally observed structural variability of chromatin in single-cells is consistent with two main co-existing sources of noise, i.e., the heterogeneity of single-cell epigenetics and, interestingly, an intrinsic conformational degeneracy, as chromatin can dynamically fold in many different conformations rather than in a single naïve structure as usual proteins.

While other folding mechanisms are likely to contribute to the organisation of the genome (such as heterochromatin adsorption to the lamina), one can speculate on why different molecular processes could cooperate in determining chromatin folding. Beyond ensuring redundancy in regulation, they appear to be more effective in implementing complementary tasks. For instance, loop-extrusion is particularly suited to establish TAD borders and pointwise strong loop interactions, whereas globule phase separation can better act to segregate different regions and to form more stable (i.e., with lower variability) and hence more reproducible regulatory structures. Additionally, while loop-extrusion requires energy consumption, phase transitions are sustained by the thermal bath, and they are robust and reversible processes as the system only needs, e.g., to set an above threshold concentration (or affinity) of binders, with no need of fine tuning their number (or strength).

## ACKNOWLEDGEMENTS

M.N. acknowledges support from the National Institutes of Health Common Fund 4D Nucleome Program grant 1U54DK107977-01 and 1UM1HG011585-01, EU H2020 Marie Curie ITN n.813282, CINECA ISCRA ID HP10CYFPS5 and HP10CRTY8P, Einstein BIH Fellowship Award (EVF-BIH-2016 and 2019), Regione Campania SATIN Project 2018-2020, and computer resources from INFN, CINECA, ENEA CRESCO/ENEAGRID (Iannone et al. 2019) and *Scope*/*ReCAS/Ibisco* at the University of Naples.

## AUTHOR CONTRIBUTIONS

M.N., M.C and E.I. designed the project. M.N., M.C., E.I. and A.M.C. developed the modelling. M.C., E.I. and A.A. ran computer simulations and performed data analyses with help from A.M.C., S.B. and A.E.. M.N. and M.C. wrote the manuscript with input from all the authors.

## REFERENCES

1. Bickmore, W. A. & Van Steensel, B. Genome architecture: Domain organization of interphase chromosomes. Cell (2013) doi:10.1016/j.cell.2013.02.001.

2. Sexton, T. & Cavalli, G. The role of chromosome domains in shaping the functional genome. Cell vol. 160 (2015).

3. Quinodoz, S. A. et al. Higher-Order Inter-chromosomal Hubs Shape 3D Genome Organization in the Nucleus. Cell 174, 744-757.e24 (2018).

4. Boettiger, A. N. et al. Super-resolution imaging reveals distinct chromatin folding for different epigenetic states. Nature 529, 418–422 (2016).

5. Cattoni, D. I. et al. Single-cell absolute contact probability detection reveals chromosomes are organized by multiple low-frequency yet specific interactions. Nat. Commun. 8, 1753 (2017).

6. Bintu, B. et al. Super-resolution chromatin tracing reveals domains and cooperative interactions in single cells. Science (80-.). (2018) doi:10.1126/science.aau1783.

7. Dekker, J. & Mirny, L. The 3D Genome as Moderator of Chromosomal Communication. Cell (2016) doi:10.1016/j.cell.2016.02.007.

8. Dixon, J. R., Gorkin, D. U. & Ren, B. Chromatin Domains: The Unit of Chromosome Organization. Mol. Cell 62, 668–680 (2016).

9. Krijger, P. H. L. & De Laat, W. Regulation of disease-associated gene expression in the 3D genome. Nature Reviews Molecular Cell Biology vol. 17 (2016).

10. Spielmann, M., Lupiáñez, D. G. & Mundlos, S. Structural variation in the 3D genome. Nat. Rev. Genet. 19, 453–467 (2018).

11. Finn, E. H. & Misteli, T. Molecular basis and biological function of variability in spatial genome organization. Science (80-.). 365, eaaw9498 (2019).

12. Kempfer, R. & Pombo, A. Methods for mapping 3D chromosome architecture. Nature Reviews Genetics vol. 21 (2020).

13. Lieberman-Aiden, E. et al. Comprehensive mapping of long-range interactions reveals folding principles of the human genome. Science (80-.). (2009) doi:10.1126/science.1181369.

14. Beagrie, R. A. et al. Complex multi-enhancer contacts captured by genome architecture mapping. Nature (2017) doi:10.1038/nature21411.

15. Rao, S. S. P. et al. A 3D map of the human genome at kilobase resolution reveals principles of chromatin looping. Cell (2014) doi:10.1016/j.cell.2014.11.021.

16. Dixon, J. R. et al. Topological domains in mammalian genomes identified by analysis of chromatin interactions. Nature (2012) doi:10.1038/nature11082.

17. Nora, E. P. et al. Spatial partitioning of the regulatory landscape of the X-inactivation centre. Nature (2012) doi:10.1038/nature11049.

18. Fraser, J. et al. Hierarchical folding and reorganization of chromosomes are linked to transcriptional changes in cellular differentiation. Mol. Syst. Biol. (2015) doi:10.15252/msb.20156492.

19. Sanborn, A. L. et al. Chromatin extrusion explains key features of loop and domain formation in wild-type and engineered genomes. Proc. Natl. Acad. Sci. U. S. A. 112, (2015).

20. Fudenberg, G. et al. Formation of Chromosomal Domains by Loop Extrusion. Cell Rep. (2016) doi:10.1016/j.celrep.2016.04.085.

21. Jost, D., Carrivain, P., Cavalli, G. & Vaillant, C. Modeling epigenome folding: Formation and dynamics of topologically associated chromatin domains. Nucleic Acids Res. (2014) doi:10.1093/nar/gku698.

22. Zhang, B. & Wolynes, P. G. Topology, structures, and energy landscapes of human chromosomes. Proc. Natl. Acad. Sci. U. S. A. 112, (2015).

23. Brackley, C. A. et al. Predicting the three-dimensional folding of cis-regulatory regions in mammalian genomes using bioinformatic data and polymer models. Genome Biol. 17, (2016).

24. Chiariello, A. M., Annunziatella, C., Bianco, S., Esposito, A. & Nicodemi, M. Polymer physics of chromosome large-scale 3D organisation. Sci. Rep. (2016) doi:10.1038/srep29775.

25. Di Stefano, M., Paulsen, J., Lien, T. G., Hovig, E. & Micheletti, C. Hi-C-constrained physical models of human chromosomes recover functionally-related properties of genome organization. Sci. Rep. 6, (2016).

26. Di Pierro, M., Zhang, B., Aiden, E. L., Wolynes, P. G. & Onuchic, J. N. Transferable model for chromosome architecture. Proc. Natl. Acad. Sci. 113, 12168–12173 (2016).

27. Barbieri, M. et al. Active and poised promoter states drive folding of the extended HoxB locus in mouse embryonic stem cells. Nat. Struct. Mol. Biol. (2017) doi:10.1038/nsmb.3402.

28. Bianco, S. et al. Polymer physics predicts the effects of structural variants on chromatin architecture. Nat. Genet. (2018) doi:10.1038/s41588-018-0098-8.

29. Buckle, A., Brackley, C. A., Boyle, S., Marenduzzo, D. & Gilbert, N. Polymer Simulations of Heteromorphic Chromatin Predict the 3D Folding of Complex Genomic Loci. Mol. Cell 72, (2018).

30. Shi, G., Liu, L., Hyeon, C. & Thirumalai, D. Interphase human chromosome exhibits out of equilibrium glassy dynamics. Nat. Commun. (2018) doi:10.1038/s41467-018-05606-6.

31. Brackley, C. A. et al. Nonequilibrium Chromosome Looping via Molecular Slip Links. Phys. Rev. Lett. 119, 138101 (2017).

32. Nuebler, J., Fudenberg, G., Imakaev, M., Abdennur, N. & Mirny, L. A. Chromatin organization by an interplay of loop extrusion and compartmental segregation. Proc. Natl. Acad. Sci. U. S. A. 115, (2018).

33. Bianco, S. et al. Modeling Single-Molecule Conformations of the HoxD Region in Mouse Embryonic Stem and Cortical Neuronal Cells. Cell Rep. (2019) doi:10.1016/j.celrep.2019.07.013.

34. Chiariello, A. M. et al. A Dynamic Folded Hairpin Conformation Is Associated with α-Globin Activation in Erythroid Cells. Cell Rep. (2020) doi:10.1016/j.celrep.2020.01.044.

35. Conte, M. et al. Polymer physics indicates chromatin folding variability across single-cells results from state degeneracy in phase separation. Nat. Commun. (2020) doi:10.1038/s41467-020-17141-4.

36. Plewczynski, D. & Kadlof, M. Computational modelling of three-dimensional genome structure. Methods vols 181–182 (2020).

37. Bianco, S. et al. Computational approaches from polymer physics to investigate chromatin folding. Current Opinion in Cell Biology (2020) doi:10.1016/j.ceb.2020.01.002.

38. Fiorillo, L. et al. Comparison of the Hi-C, GAM and SPRITE methods using polymer models of chromatin. Nat. Methods 18, (2021).

39. Racko, D., Benedetti, F., Dorier, J. & Stasiak, A. Transcription-induced supercoiling as the driving force of chromatin loop extrusion during formation of TADs in interphase chromosomes. Nucleic Acids Res. 46, (2018).

40. Banigan, E. J. & Mirny, L. A. Loop extrusion: theory meets single-molecule experiments. Current Opinion in Cell Biology vol. 64 (2020).

41. Nicodemi, M. & Pombo, A. Models of chromosome structure. Current Opinion in Cell Biology (2014) doi:10.1016/j.ceb.2014.04.004.

42. Nicodemi, M. & Prisco, A. Thermodynamic pathways to genome spatial organization in the cell nucleus. Biophys. J. (2009) doi:10.1016/j.bpj.2008.12.3919.

43. Bohn, M. & Heermann, D. W. Diffusion-driven looping provides a consistent provides a consistent framework for chromatin organization. PLoS One (2010) doi:10.1371/journal.pone.0012218.

44. Barbieri, M. et al. Complexity of chromatin folding is captured by the strings and binders switch model. Proc. Natl. Acad. Sci. (2012) doi:10.1073/pnas.1204799109.

45. Brackley, C. A., Taylor, S., Papantonis, A., Cook, P. R. & Marenduzzo, D. Nonspecific bridging-induced attraction drives clustering of DNA-binding proteins and genome organization. Proc. Natl. Acad. Sci. U. S. A. (2013) doi:10.1073/pnas.1302950110.

46. Lesne, A., Riposo, J., Roger, P., Cournac, A. & Mozziconacci, J. 3D genome reconstruction from chromosomal contacts. Nat. Methods 11, (2014).

47. Tjong, H. et al. Population-based 3D genome structure analysis reveals driving forces in spatial genome organization. Proc. Natl. Acad. Sci. U. S. A. (2016) doi:10.1073/pnas.1512577113.

48. Zhang, S., Chasman, D., Knaack, S. & Roy, S. In silico prediction of high-resolution Hi-C interaction matrices. Nat. Commun. 10, (2019).

49. Fudenberg, G., Kelley, D. R. & Pollard, K. S. Predicting 3D genome folding from DNA sequence with Akita. Nat. Methods 17, (2020).

50. Schwessinger, R. et al. DeepC: predicting 3D genome folding using megabase-scale transfer learning. Nat. Methods 17, (2020).

51. Wang, Y. et al. SPIN reveals genome-wide landscape of nuclear compartmentalization. Genome Biol. 22, (2021).

52. Li, Q. et al. The three-dimensional genome organization of Drosophila melanogaster through data integration. Genome Biol. 18, 145 (2017).

53. Serra, F. et al. Automatic analysis and 3D-modelling of Hi-C data using TADbit reveals structural features of the fly chromatin colors. PLoS Comput. Biol. 13, (2017).

54. Nir, G. et al. Walking along chromosomes with super-resolution imaging, contact maps, and integrative modeling. PLoS Genet. 14, e1007872 (2018).

55. Lin, D., Bonora, G., Yardimci, G. G. & Noble, W. S. Computational methods for analyzing and modeling genome structure and organization. Wiley Interdiscip. Rev. Syst. Biol. Med. 11, e1435 (2018).

56. Di Stefano, M., Paulsen, J., Jost, D. & Marti-Renom, M. A. 4D nucleome modeling. Current Opinion in Genetics and Development vol. 67 (2021).

57. Kim, H. J. et al. Capturing cell type-specific chromatin compartment patterns by applying topic modeling to single-cell Hi-C data. PLoS Comput. Biol. 16, (2020).

58. Marti-Renom, M. A. Benchmarking experiments with polymer modeling. Nature Methods vol. 18 (2021).

59. Qi, Y. & Zhang, B. Predicting three-dimensional genome organization with chromatin states. PLoS Comput. Biol. 15, (2019).

60. Rao, S. S. P. et al. Cohesin Loss Eliminates All Loop Domains. Cell 171, 305-320.e24 (2017).

61. Alberts, B. et al. Molecular Biology of the Cell. Molecular Biology of the Cell (2007). doi:10.1201/9780203833445.

62. De Gennes, P. G. Scaling concepts in polymer physics. Cornell university press. Ithaca N.Y., (1979) doi:10.1163/_q3_SIM_00374.

63. Grant, C. E., Bailey, T. L. & Noble, W. S. FIMO: Scanning for occurrences of a given motif. Bioinformatics 27, (2011).

64. Dunham, I. et al. An integrated encyclopedia of DNA elements in the human genome. Nature 489, 57–74 (2012).

65. Carter, B. & Zhao, K. The epigenetic basis of cellular heterogeneity. Nature Reviews Genetics vol. 22 (2021).

66. Stevens, T. J. et al. 3D structures of individual mammalian genomes studied by single- cell Hi-C. Nature 544, 59–64 (2017).

67. Huang, H. et al. CTCF mediates dosage- and sequence-context-dependent transcriptional insulation by forming local chromatin domains. Nat. Genet. 53, (2021).

68. Kremer, K. & Grest, G. S. Dynamics of entangled linear polymer melts: A molecular- dynamics simulation. J. Chem. Phys. (1990) doi:10.1063/1.458541.

69. Plimpton, S. Fast parallel algorithms for short-range molecular dynamics. J. Comput. Phys. (1995) doi:10.1006/jcph.1995.1039.

70. Anderson, J. A., Glaser, J. & Glotzer, S. C. HOOMD-blue: A Python package for high- performance molecular dynamics and hard particle Monte Carlo simulations. Comput. Mater. Sci. 173, (2020).

71. Goloborodko, A., Marko, J. F. & Mirny, L. A. Chromosome Compaction by Active Loop Extrusion. Biophys. J. 110, (2016).

72. Rosa, A. & Everaers, R. Structure and dynamics of interphase chromosomes. PLoS Comput. Biol. (2008) doi:10.1371/journal.pcbi.1000153.

73. Allen, M. P. & Tildesley, D. J. Computer Simulation of Liquids (Oxford Science Publications) SE - Oxford science publications. Oxford Univ. Press (1989).

74. Theobald, D. L. Rapid calculation of RMSDs using a quaternion-based characteristic polynomial. Acta Crystallogr. Sect. A Found. Crystallogr. 61, (2005).

