## Supplementary Material for "Loop-extrusion and polymer phase-separation can co-exist at the single-molecule level to shape chromatin folding"

### for

#### SUPPLEMENTARY FIGURES

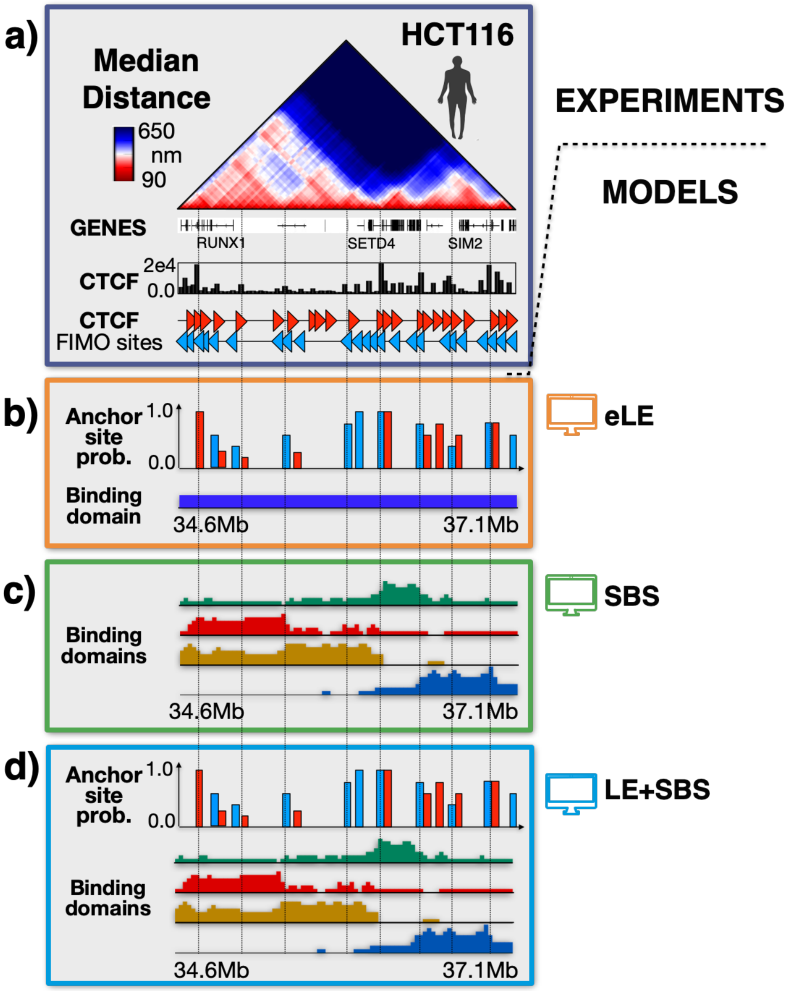


**Supplementary Figure 1. Scheme of the investigated polymer models in HCT116.**

**a)** Microscopy median distance[1] and ENCODE[2] CTCF data are shown for the studied 2.5Mb wide locus in human HCT116 cells. **b)** We considered an extended LE (eLE) model where the genomic locations of the anchor sites are optimized, independently of CTCF, to best reproduce Hi-C and average microscopy data. Also, to mimic single-cell epigenetic heterogeneity, those anchor sites have a specific, finite probability to be present in a model single molecule[3]. **c)** The Strings and Binders (SBS) model[4] of the locus has four distinct types of binding sites (represented by different colors), inferred by the PRISMR method and each correlated with different combinations of epigenetic factors including, but not limited to, CTCF and cohesin[4,5]. **d)** Scheme of the combined LE+SBS polymer model where both the eLE and SBS mechanisms act simultaneously in each single-molecule conformation.

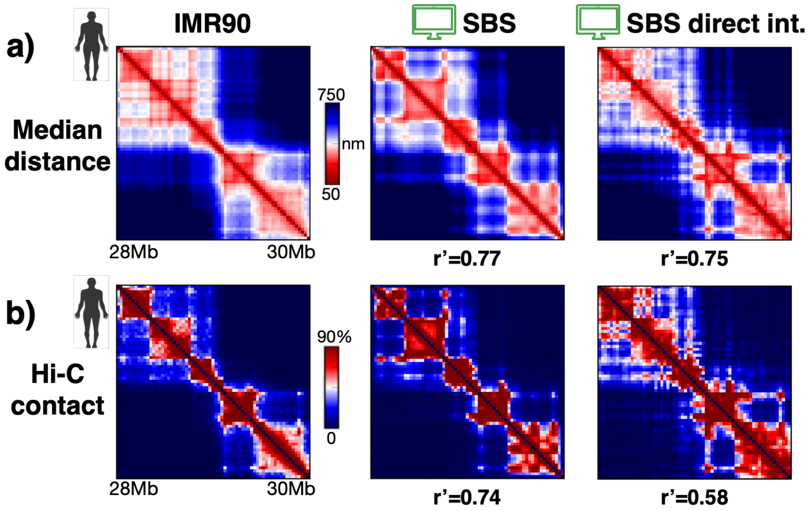


**Supplementary Figure 2. A variant of the SBS model with direct interactions between DNA sites, rather than mediated by molecular binders, has behaviors analogous to the original SBS.**

We explored a variant of the SBS model where cognate DNA sites have direct physical interactions, rather than mediated by diffusing binders. **a)** The median distance matrix[1] and **b)** the Hi-C contact map[6] of the IMR90 locus (left) are well recapitulated by both the SBS model (middle) and its variant with direct physical interactions (right), as quantified by their high genomic distance-corrected Pearson correlations (r') with the experiments. In particular, the model with direct interactions returns contact patterns similar to the SBS, as expected from Statistical Mechanics[7].

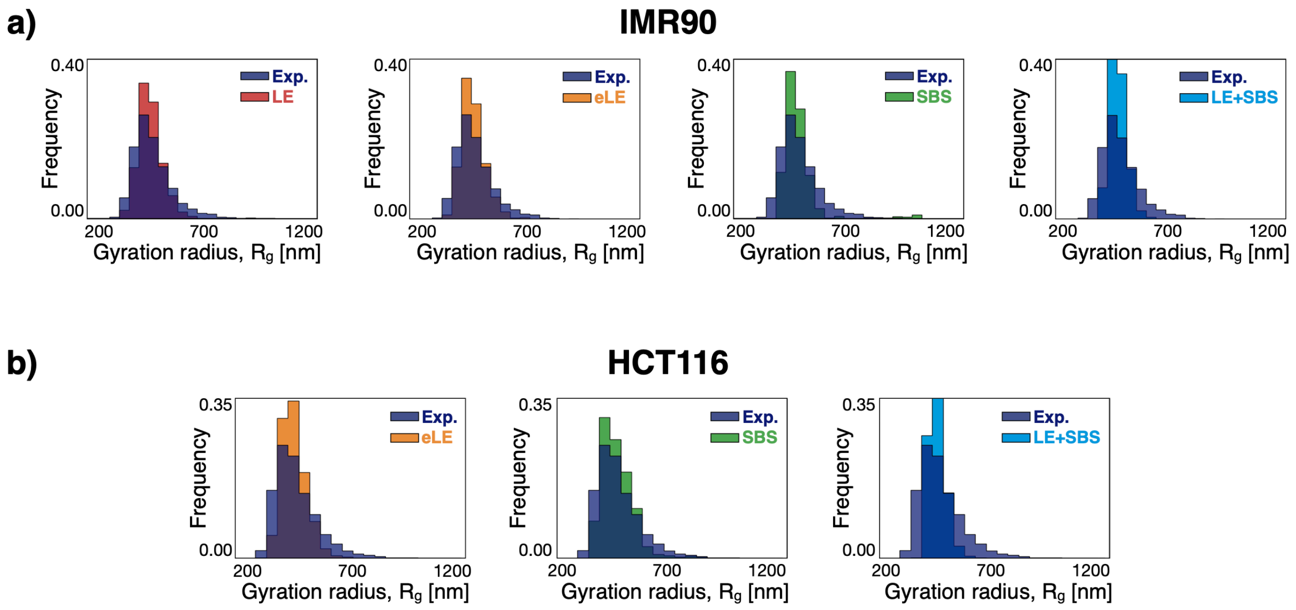


**Supplementary Figure 3. Gyration radius distributions in imaging experiments and in the models of the studied loci.**

Experimental[1] and model derived gyration radius distributions in **a)** IMR90 and **b)** HCT116 cells. In all the considered cases, the model unit length scale is mapped into physical units by equating the median gyration radius to its corresponding experimental value.

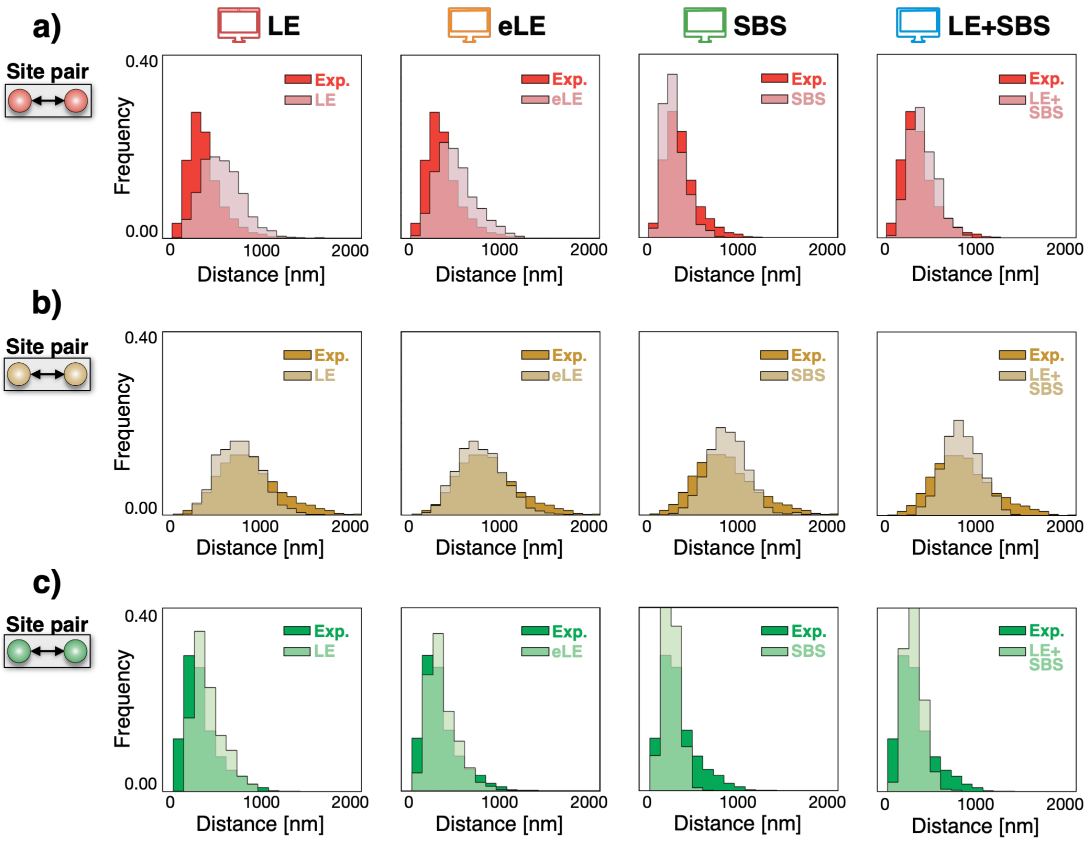


**Supplementary Figure 4. Distance distributions of site pairs in the IMR90 locus.**

We measured in both the experiment[1] and models the distance distributions of specific, interesting pairs of sites along the IMR90 locus. Distributions are shown for: **a)** a pair of sites (red), located 0.7Mb away in different sub-TADs and forming a strong loop contact in bulk data; **b**) a pair of sites (yellow) 1.1Mb apart from different TADs and separated by a strong TAD boundary in between; **c)** a pair (green) of 0.3Mb distant sites with a strong loop interaction in the same TAD. The distributions derived from the models are all similar to the corresponding imaged distributions, yet the specific values of those distances can depend on the minute details of the models.

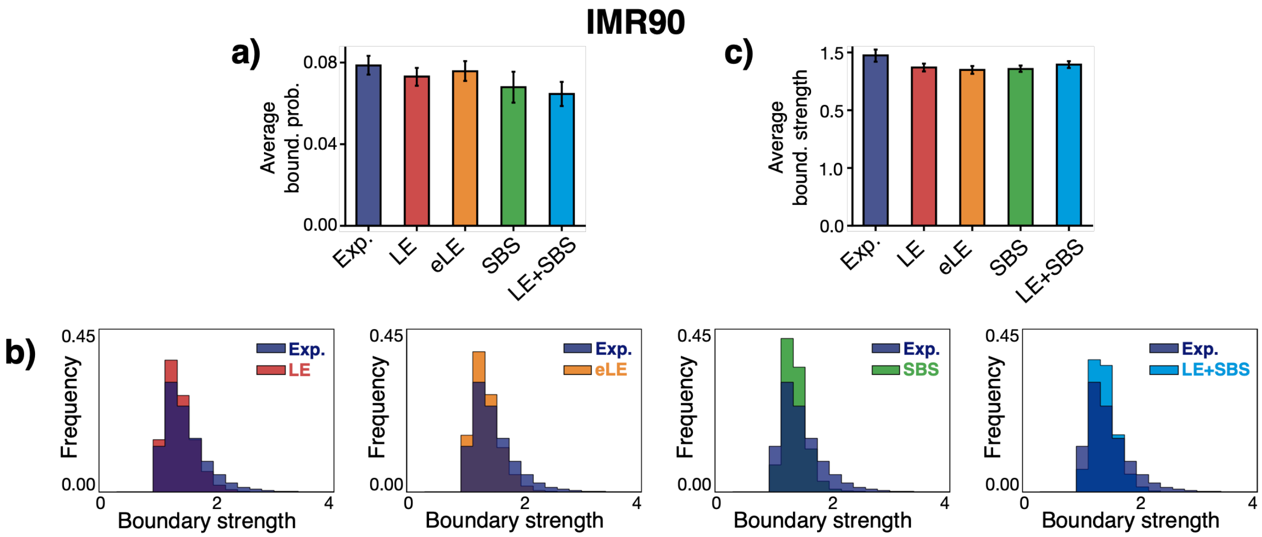


**Supplementary Figure 5. The average boundary probability and strength are similar across the different models and close to the corresponding experimental values in IMR90.**

**a)** TAD boundary probability averaged over all genomic positions in both experiment[1] and models in the IMR90 locus (error bars s.e.m.). **b)** The distributions of the boundary strengths derived from the models are all similar to the experimental distribution, as well as **c)** their corresponding average values (error bars s.e.m.).

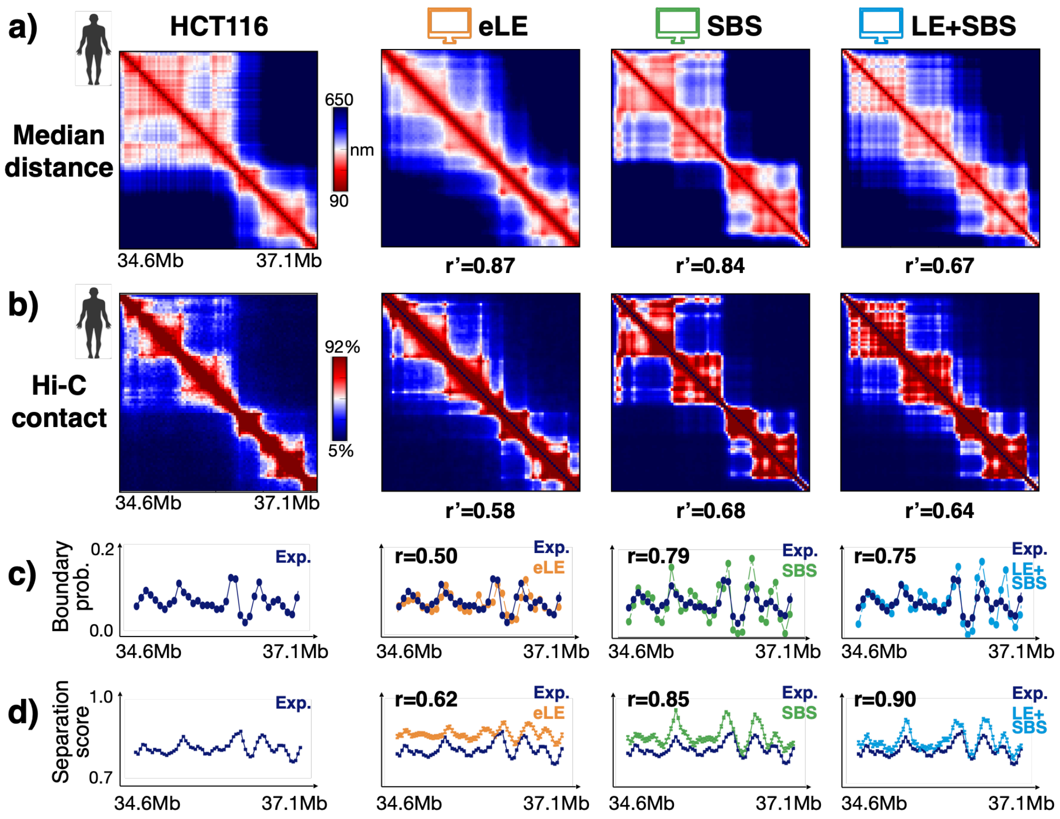


**Supplementary Figure 6. Both loop-extrusion and phase-separation based models recapitulate bulk Hi-C and average microscopy data in the HCT116 locus.**

**a)** In-silico median distance and **b)** average contact data are compared to microscopy[1] and Hi-C[8] data (left) in the HCT116 locus. The different models have all high genomic distance-corrected Pearson correlations, r', with the experiments. **c)** The average single-molecule genomic boundary probability and **d)** the separation score are also well recapitulated by the models.

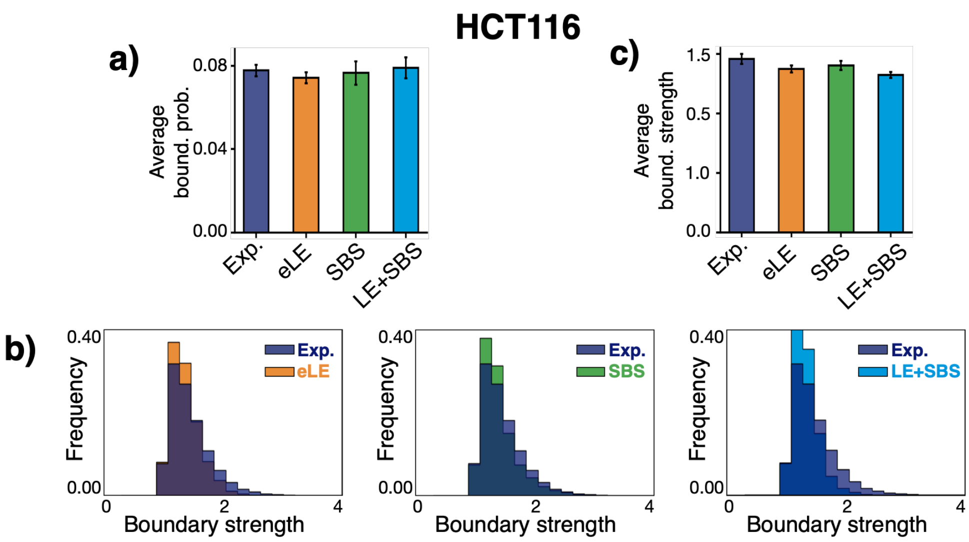


**Supplementary Figure 7. The average boundary probability and strength are similar across the different models and close to the corresponding experimental values in HCT116.**

**a)** TAD boundary probability averaged over all genomic positions in both experiment[1] and models in the HCT116 locus (error bars s.e.m.). **b)** The distributions of the boundary strengths derived from the models are all similar to the experimental distribution, as well as **c)** their corresponding average values (error bars s.e.m.).

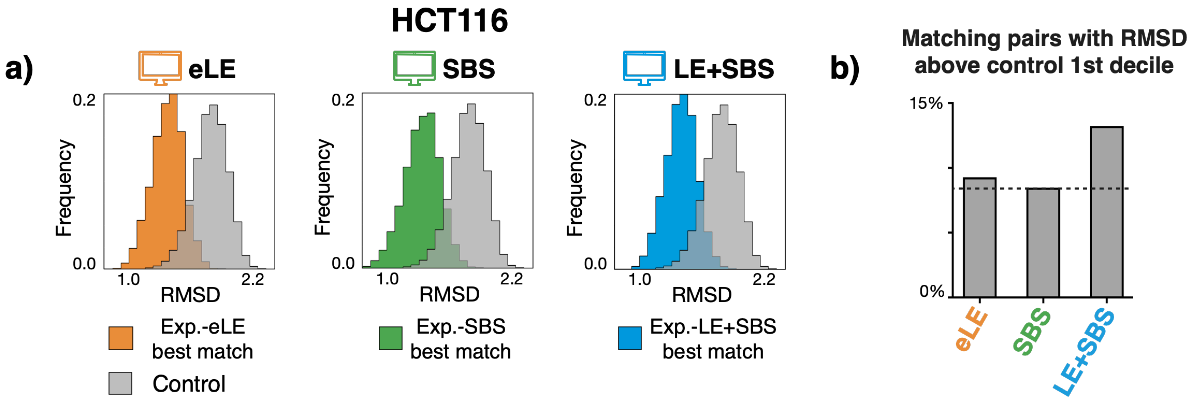


**Supplementary Figure 8. The best-matching experiment-model pairs RMSD distributions are statistically different from a random control in HCT116.**

**a)** For each of the considered polymer models, the RMSD distribution of the best-matching experiment-model pairs in HCT116 is statistically different from a control RMSD distribution made of random pairs of experimental structures[1] (two-sided Mann–Whitney test p-value = 0). **b)** Less than 15% of the best matching pairs have an RMSD above the 1st decile of the control distribution. The SBS, in particular, performs slightly better than the other models.

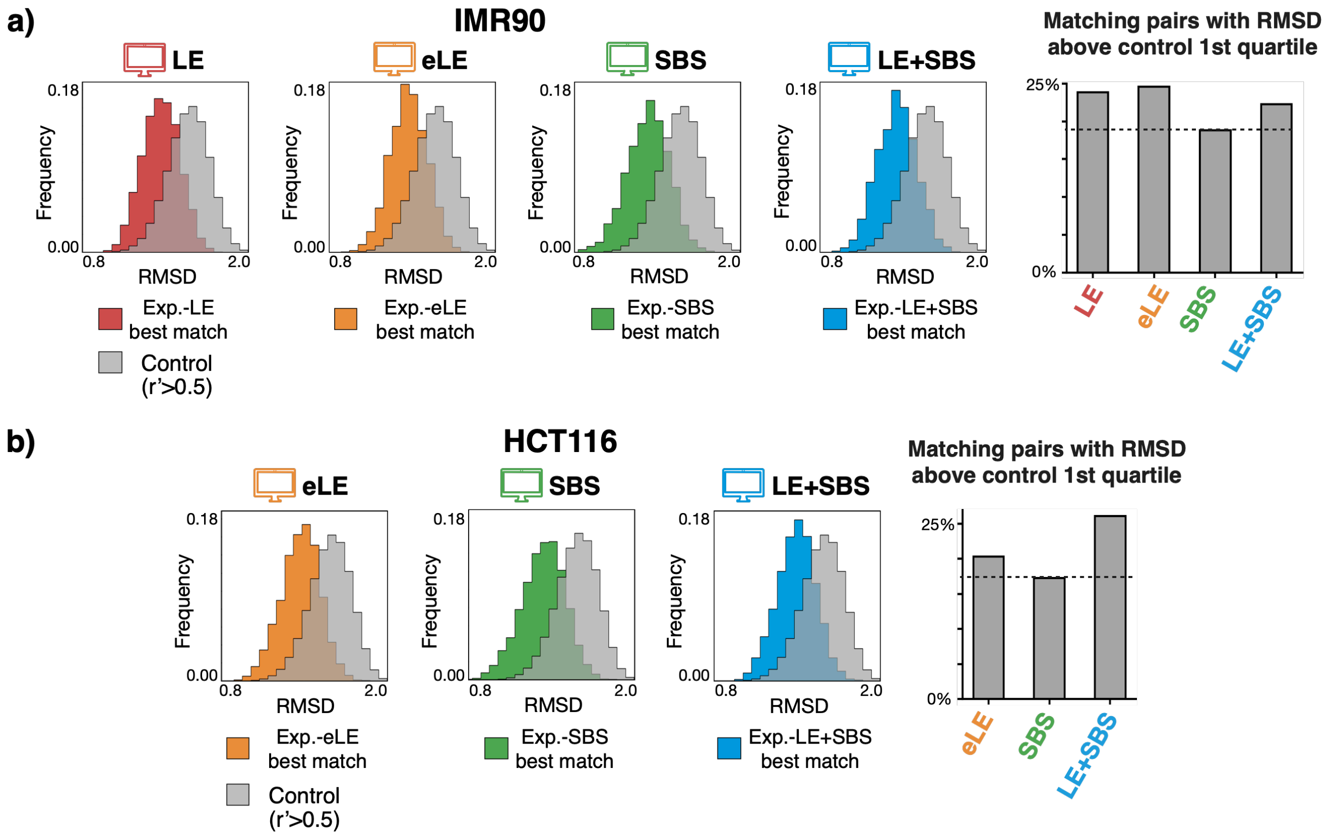


**Supplementary Figure 9. An additional, more stringent control to test the statistical significance of the RMSD association.**

To test the significance of the best-matching experiment-model pairs identified via the RMSD criterion, we performed an additional, more stringent control where the RMSD is computed only between pairs of microscopy conformations[1] having distance matrices with a corresponding genomic distance-corrected Pearson correlation value r'>0.5. We found that in both the **a)** IMR90 and **b)** HCT116 loci the RMSD distributions of the best-matches of each model are statistically different from the control (left, two-sided Mann–Whitney test p-value = 0) with less than 25% of entries of the former falling above the 1st quartile of the latter (right). Hence, the model 3D conformations best matching the imaged structures have a statistically significant RMSD distribution and provide a non-trivial description of chromatin structure in single molecules.

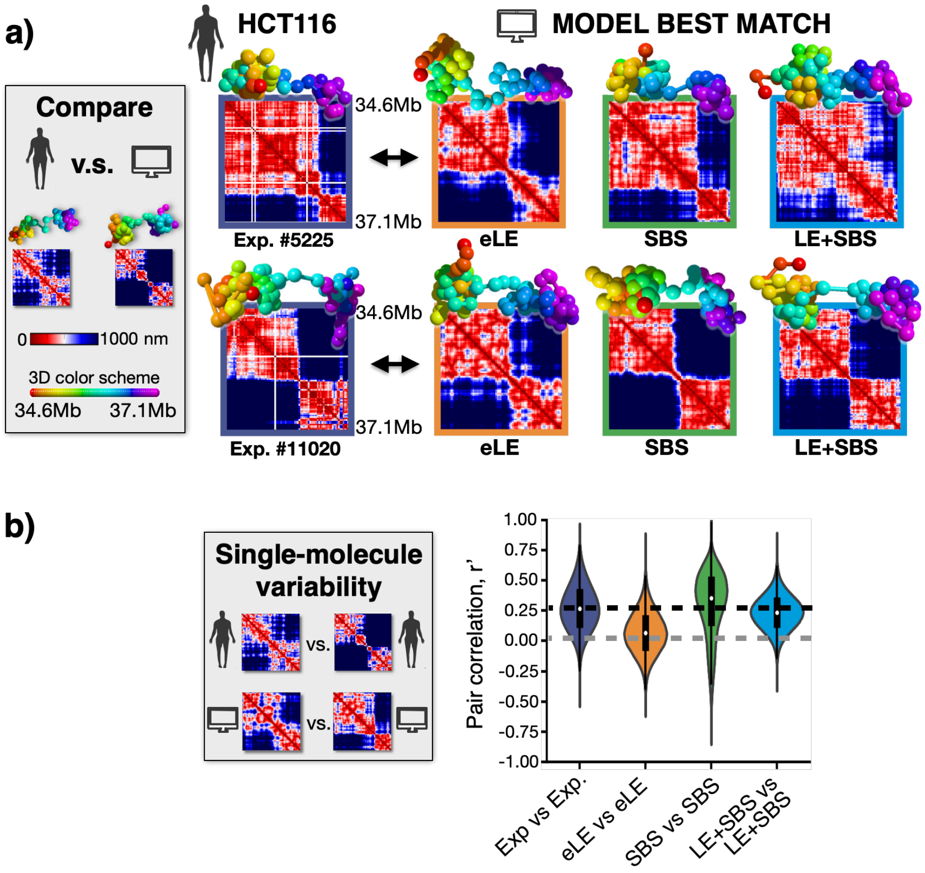


**Supplementary Figure 10. Single-cell chromatin conformations of the HCT116 locus are well captured by the models, especially by phase-separation based ones.**

**a)** Microscopy single-cell chromatin structures of the HCT116 locus[1] (left) are associated to a best matching single-molecule conformation in each model via the minimum RMSD criterion. Two examples are shown here. **b**) The variability of single-cell imaged structures is measured by the distribution of r' correlations between pairs of distance matrices and is compared to the variability of in-silico structures. The experimental distribution is broad and has an average r'=0.27, while the corresponding average values from the models are: r'=0.07, r'=0.30, and r'=0.23, respectively, for the eLE, SBS, and LE+SBS.

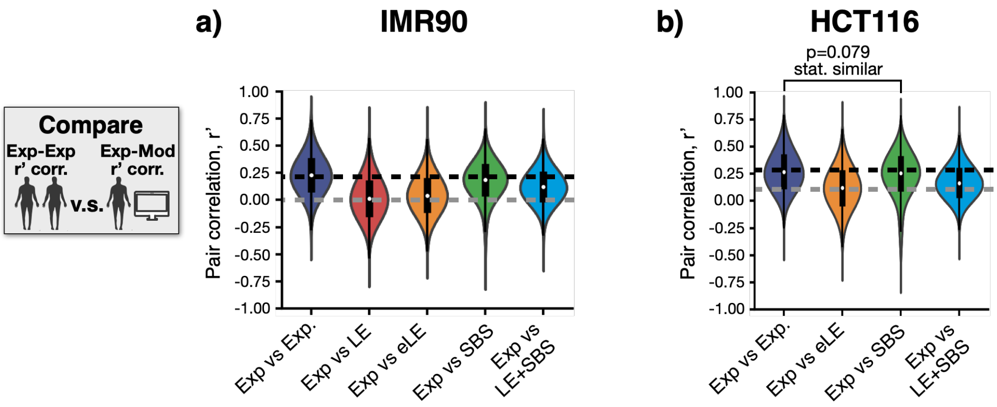


**Supplementary Figure 11. Distributions of r' correlations between experiment and model single-molecule distance matrices.**

**a)** The distribution of r' correlations between pairs of single-cell microscopy distance matrices[1] is compared, for each of the considered models, to the experiment-model r' distribution in the case of the IMR90 locus. **b)** Same as in panel **a)** for the HCT116 locus. In particular, the r' distribution in the case of the SBS model is statistically indistinguishable from the experimental one (two-sided Mann–Whitney test p-value = 0.079).

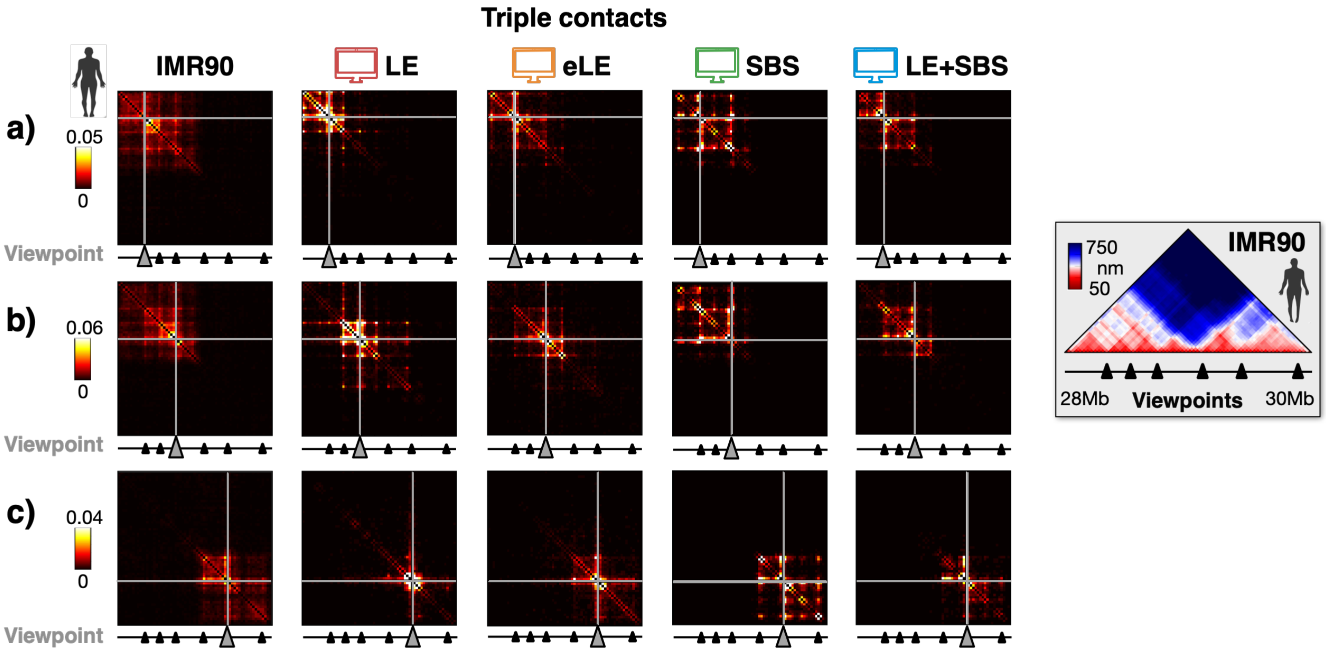


**Supplementary Figure 12. Triple contact data from additional viewpoints in IMR90.**

Triple contact probability maps are shown in microscopy data[1] (left) and in the models from three different, additional viewpoints in the IMR90 locus (panels **a)**, **b)**, **c)**). The experimental triplet contact patterns are well captured by the different models, especially by the eLE, SBS and LE+SBS.

#### SUPPLEMENTARY TABLES

**a) IMR90**

Pearson correlation r
Model vs Exp.

| | Median distance | Hi-C contact |
|---|---|---|
| **LE** | r=0.90 | r=0.87 |
| **eLE** | r=0.93 | r=0.95 |
| **SBS** | r=0.96 | r=0.94 |
| **LE+SBS** | r=0.95 | r=0.94 |

**b) HCT116**

Pearson correlation r
Model vs Exp.

| | Median distance | Hi-C contact |
|---|---|---|
| **eLE** | r=0.94 | r=0.97 |
| **SBS** | r=0.95 | r=0.88 |
| **LE+SBS** | r=0.92 | r=0.89 |

**Supplementary Table I.**

Pearson correlation, r, values between model-derived and experimental median distance[1] and Hi-C[6,8] contact maps of the studied **a)** IMR90 and **b)** HCT116 loci.

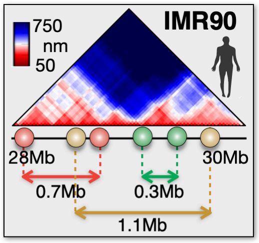


| | a) Av. dist.+-st.d [nm] | b) Av. dist.+-st.d [nm] | c) Av. dist.+-st.d [nm] |
|---|---|---|---|
| | 370+-190 | 860+-330 | 340+-200 |
| LE | 560+-220 | 750+-240 | 410+-160 |
| eLE | 510+-220 | 780+-260 | 360+-160 |
| SBS | 300+-130 | 880+-220 | 260+-90 |
| LE+SBS | 410+-150 | 840+-210 | 310+-100 |

**Supplementary Table II.**

Average and standard deviation (in nm) of the physical distances of the IMR90 site pairs investigated in **Supplementary Fig. 4**.

**REFERENCES**


1. Bintu, B. *et al.* Super-resolution chromatin tracing reveals domains and cooperative interactions in single cells. *Science (80-. ).* (2018) doi:10.1126/science.aau1783.

2. Dunham, I. *et al.* An integrated encyclopedia of DNA elements in the human genome. *Nature* **489**, 57–74 (2012).

3. Buckle, A., Brackley, C. A., Boyle, S., Marenduzzo, D. & Gilbert, N. Polymer Simulations of Heteromorphic Chromatin Predict the 3D Folding of Complex Genomic Loci. *Mol. Cell* **72**, (2018).

4. Conte, M. *et al.* Polymer physics indicates chromatin folding variability across single-cells results from state degeneracy in phase separation. *Nat. Commun.* (2020) doi:10.1038/s41467-020-17141-4.

5. Bianco, S. *et al.* Polymer physics predicts the effects of structural variants on chromatin architecture. *Nat. Genet.* (2018) doi:10.1038/s41588-018-0098-8.

6. Rao, S. S. P. *et al.* A 3D map of the human genome at kilobase resolution reveals principles of chromatin looping. *Cell* (2014) doi:10.1016/j.cell.2014.11.021.

7. De Gennes, P. G. Scaling concepts in polymer physics. Cornell university press. *Ithaca N.Y.,* (1979) doi:10.1163/_q3_SIM_00374.

8. Rao, S. S. P. *et al.* Cohesin Loss Eliminates All Loop Domains. *Cell* **171**, 305-320.e24 (2017).
